# CDC14 phosphatases control adipogenesis via PPARγ de-phosphorylation

**DOI:** 10.64898/2025.12.13.694119

**Authors:** Diana Vara-Ciruelos, A. Filipa Martins, Aicha El-Bakkali, Agustín Sánchez-Belmonte, Osvaldo Graña-Castro, Eduardo Zarzuela, Marta Isasa, María Salazar-Roa, Carolina Villarroya-Beltri, Marcos Malumbres

## Abstract

Adipogenesis is a finely tuned process requiring an appropriate balance between proliferation and differentiation. We demonstrate that CDC14, a CDK-counteracting phosphatase, regulates adipogenesis and glucose metabolism in mammals. Simultaneous depletion of CDC14A and CDC14B protects against high-fat diet-induced obesity, hepatic steatosis and glucose intolerance *in vivo*. Lack of these phosphatases blocks differentiation of stem cells into adipocytes, a defect rescued by the PPARγ agonist troglitazone. Mechanistically, we show that CDC14 counteracts ERK2 and CDK5-induced phosphorylation of PPARγ, a master regulator of adipogenesis, by directly dephosphorylating S112, S273 and T296 residues. This study supports the idea that cell cycle regulators in multicellular organisms evolved to coordinate the balance between proliferation and differentiation in different cell types. Furthermore, since PPARγ is a key target of anti-diabetic drugs and a central regulator of adipogenesis and metabolism, these findings may have important implications for treatment of metabolic diseases such as type 2 diabetes.

## Introduction

Protein phosphorylation is one of the most important posttranslational regulatory mechanisms in cells, being involved in the control of virtually every cellular process. The cell cycle is the paradigmatic example of the importance of phosphorylation in the regulation of cellular functions, as it is exquisitely coordinated by a succession of phosphorylation events carried out by cyclin-dependent kinases (CDKs) among other kinase families^1^. Although traditionally much more attention has been given to the role of kinases, phosphatases are also essential regulators of the cell cycle^2^. The dual-specificity CDC14 phosphatase is the main CDK-counteracting phosphatase in *Saccharomyces cerevisiae*, thereby allowing ordered progression through the cell cycle and eventual exit from it^3^. However, in mammals this function has been mostly assumed by PP2A complexes, and the role of CDC14 phosphatases has remained obscure due to contradictory results and possible compensatory effects between the different *CDC14* genes^4^. Hence, different *in vitro* studies have linked CDC14A to centrosome separation and spindle formation^5,6^, whereas CDC14B has been involved in cell cycle progression, nuclear organization, centriole duplication, mitotic spindle assembly, transcriptional activation and DNA damage^7–11^. In a recent work, we generated *Cdc14a* and *b* double knockout mice, and proposed a role for these phosphatases in mitosis exit in progenitor cells and differentiation^12^. This work uncovered an unexpected role of CDC14 phosphatases in the control of stem cell differentiation into neurons by de-phosphorylating the epigenetic regulator UTF1^12^. Since CDKs maintain pluripotency through the regulation of several stemness factors like OCT4, SOX2, or NANOG, epigenetic factors or the PI3K/AKT pathway^13–16^, these data support a model in which the control of phosphorylation by the CDK-CDC14 balance may control exit from the progenitor cell cycle to favor differentiation or maturation towards specific cell lineages.

The coordination of proliferation and differentiation processes is particularly crucial for adipogenesis, during which adipocyte progenitors first reenter in the cell cycle and undergo a phase of proliferation, followed by a second phase of cell cycle arrest and terminal differentiation into adipocytes. Throughout these transitions, there is significant crosstalk between proliferation and differentiation pathways, with various cell cycle regulators playing key roles. For example, E2F1 positively regulates the expression of peroxisome proliferator-activated receptor γ (PPARγ)^17^, which is the master regulator of adipocyte differentiation, whereas RB1 inhibits PPARγ activity through direct protein-protein interaction^18^. In addition, CDK4 directly activates PPARγ thereby promoting adipogenesis^19^. However, CDK4 is not the only CDK involved in the regulation of PPARγ, other CDK family members such as CDK5, CDK7 and CDK9 can also regulate PPARγ activity through direct phosphorylation at different residues^20^. CDK5 has been shown to directly phosphorylate PPARγ at S273, leading to decreased insulin-sensitizing activities while preserving its adipogenic function^21^. In contrast, CDK7 and CDK9, along with ERK kinases, phosphorylate S112 affecting the transcriptional activity of PPARγ and thereby altering its adipogenic potential. The strict regulation of PPARγ is crucial, given the profound impact of adipogenesis on metabolic health. Notably, thiazolidinediones—antidiabetic drugs that enhance insulin sensitivity in patients with type 2 diabetes—act as PPARγ agonists and can influence its phosphorylation status^21^. Therefore, understanding the factors that modulate PPARγ phosphorylation holds significant therapeutic potential for diabetes management. In this study, we reveal that CDC14 phosphatases directly dephosphorylate PPARγ, thereby regulating adipogenesis and insulin sensitivity *in vivo*.

## Results

### Deficient adipose tissue formation in *Cdc14a; Cdc14b* double knockout mice

Previous data from our laboratory showed that *Cdc14a*; *Cdc14b* double KO (*Cdc14* DKO) mice display a mortality rate of approximately 50% within the first 3 weeks of life, accompanied by defects in brain development^12^. *Cdc14* DKO mice that survived the perinatal period, developed normally, were fertile and did not exhibit major abnormalities or disorders, as concluded from a general histological analysis of tissues at 8 and 15 weeks (data not shown). In addition, *Cdc14* DKO mice exhibited a mortality rate similar to wild-type mice throughout a period of 90 weeks without increased aged-related pathologies (Figure 1A). However, *Cdc14* DKO pups were also characterized by a striking paucity of adipose tissue concomitant with defective accumulation of fat in adipocytes (Figure 1B). In fact, *Cdc14* DKO mice displayed reduced weight at early ages (Figure 1C) and 90 weeks of age (Figure 1D). Detailed histological evaluation of the tissue samples obtained from these mice revealed decreased incidence of hepatic steatosis of *Cdc14* DKO when compared to wild type mice (Figure 1E). Furthermore, *Cdc14* DKO mice have smaller adipocyte cell size in white adipose tissue (WAT) (Figure 1F) and reduced lipid accumulation in brown adipose tissue (BAT) (Figure 1G).

**Figure 1.**
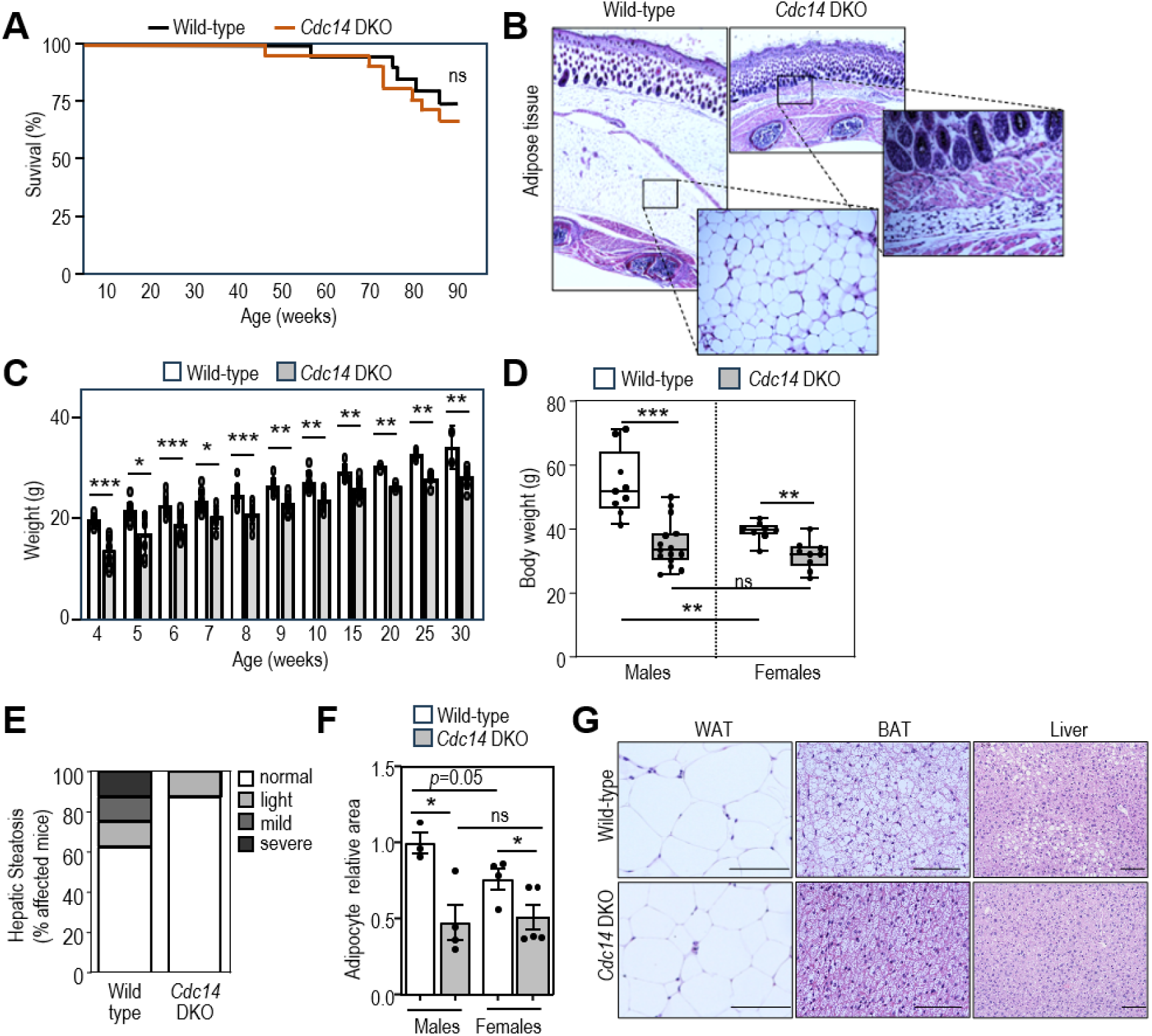
*Cdc14* DKO adult mice exhibit reduced body weight and adipose disfunction. **A**. Kaplan-Mayer curve of the survival of control (n=19) and double-deficient Cdc14a; Cdc14b (Cdc14 DKO; n=21) mice during the first 90 weeks. **B**. Hematoxylin and eosin (H&E) images of adipose tissue of wild type and *Cdc14* DKO mice at day 9 after birth. **C**. Body weight of control mice and *Cdc14* DKO mice that survived preweaning lethality at the indicated time points. Each bar represents a minimum of 3 animals per each phenotype. **D**. Body weight of males (left) and females (right) of wild type and *Cdc14* DKO mice at 90 weeks of age. N=9 male wild type, N=15 male *Cdc14* DKO, N=9 female wild type, N=9 female *Cdc14* DKO. **E**. Incidence of hepatic steatosis in wild type and *Cdc14* DKO mice at 90 weeks of age. N=8 wild type, N=8 *Cdc14* DKO. **F**. Quantification of the area of the adipocyte cells that composed the epididymal WAT of wild type and *Cdc14* DKO mice at 90 weeks of age. Each point represents the average relative size area of at least 50 adipocyte cells of each individual mice. N=3 male wild type, N=4 male *Cdc14* DKO, N=4 female wild type, N=5 female *Cdc14* DKO. **G**. Representative H&E images of white adipose tissue (WAT), brown adipose tissue (BAT) and liver sections of wild type and *Cdc14* DKO mice at 90 weeks of age. Scale bars, 100 μm. Data are shown as mean ±SEM. Statistical significance was assessed using the two-tailed Student’s t-test with Welch correction for all data with the exception of A., where the Chi-square test was applied. ns, not significant (*p*>0.05); *, *p*<0.05; **, *p*<0.005.

### Differential effects of CDC14A and CDC14B in diet-induced obesity

To determine the contribution of CDC14A and CDC14B to adipocyte differentiation and fat accumulation, we subjected *Cdc14a* and *Cdc14b* single knockout mice to high-fat diet (HFD) conditions. Single *Cdc14b* KO mice gained weight with similar kinetics to that wild-type controls (Figure 2A and Supplementary Figure 1A,B), and no significant differences were found in hepatic steatosis incidence, WAT accumulation or adipocyte area (Figure 2B,C,D). White fat deposition into brown fat and pancreatic Langerhans islets size were also similar between the two groups (Figure 2E,F).

**Figure 2.**
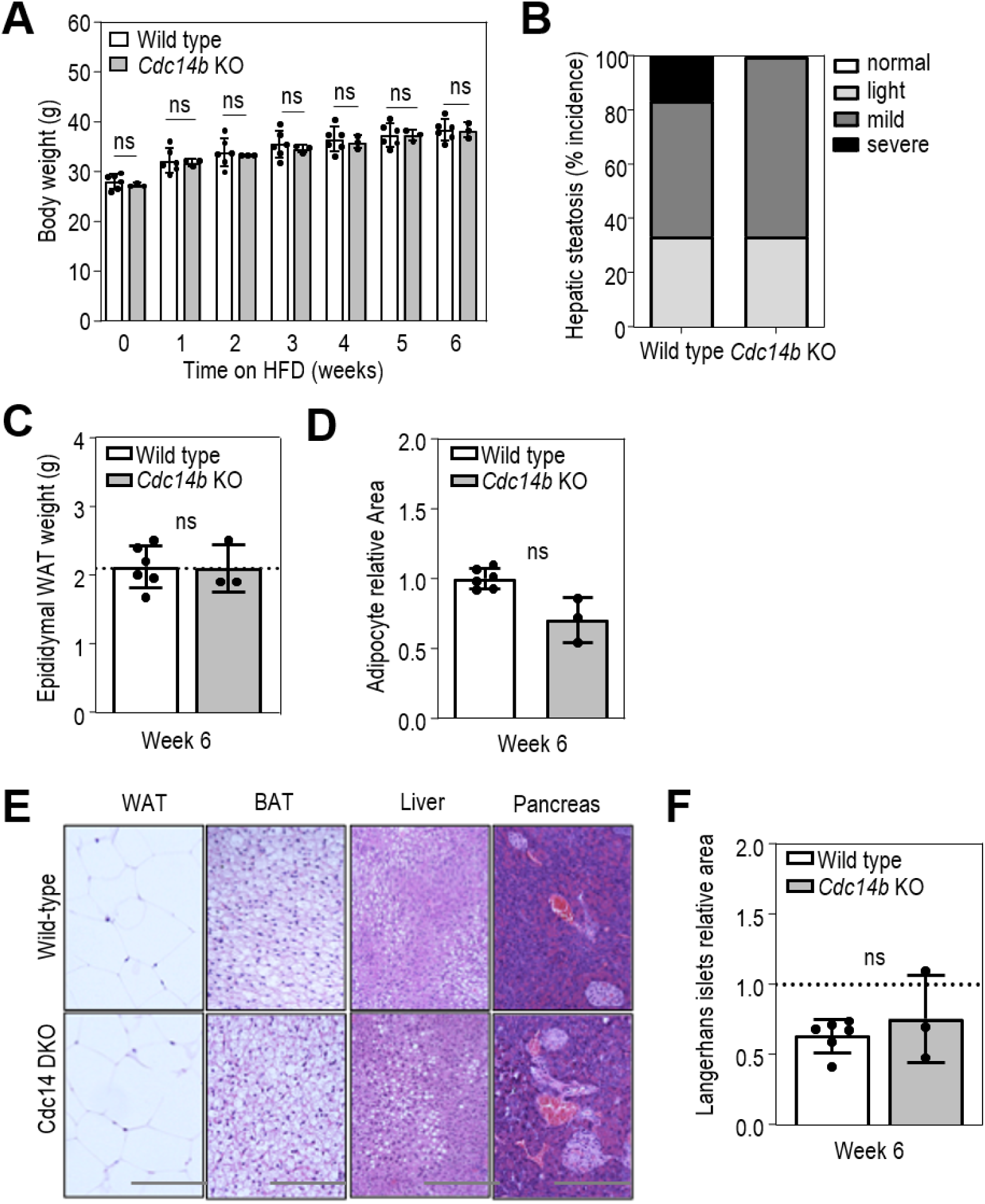
*Cdc14b* KO display normal physiological response to high fat diet. **A**. Body weight of wild type and *Cdc14b* KO mice after 6 weeks of high fat diet (HFD). **B**. Incidence of hepatic steatosis in wild type and *Cdc14b* KO mice at the end of the HFD treatment. **C**. Weight of the epidydimal WAT of wild type and *Cdc14b* KO mice after the HFD treatment. **D**. Quantification of the average area of the adipocyte cells that composed the WAT in wild type and *Cdc14* DKO mice after HFD. Each point represents the average relative size area of at least 50 adipocytes of each mouse. **E**. Representative H&E staining of WAT, BAT, liver and pancreatic tissue sections of wild type and *Cdc14b* KO mice at the end of HFD treatment. N=6 wild type, N=3 Cdc14b KO for HFD treatment. All mice used in this experimental setting were male between 15 and 18 weeks of age. Scale bars, 100 μm. **F**, Quantification of average area of pancreatic B-islets in wild type and Cdc14b KO mice after 6 weeks of HFD. Each dot represents the average relative area of at least 5 B-islets per mice. Data are shown as mean ±SEM. Statistical significance was assessed using 2-way ANOVA (Bonferroni’s multiple test) for A, B and C, and the two-tailed unpaired Student’s t-test with Welch correction for E. ns, not significant (p>0.05).

Unexpectedly, single *Cdc14a* KO mice gained significantly more weight than wild-type mice (Figure 3A). After 6 weeks of treatment, *Cdc14a* KO mice showed a cumulative weight gain of 60%, twice the value of control mice (Supplementary Figure 1C,D). Whereas the increase in weight dropped rapidly in wild-type mice, stabilizing at week 4 with a value around 2.6% of mice body weight, *Cdc14a* KO weight variation actually increased significantly by the second week and displayed a general tendency to be higher than control values at all-time points until the end of the experiment (Supplementary Figure 1D). The incidence and severity of hepatic steatosis observed in the *Cdc14a* KO experimental group was higher as well, with 100% of these mice affected by severe steatosis (Figure 3B). The exacerbation of the phenotype was also evident at the level of WAT accumulation (Figure 3C), the extent of adipocyte hypertrophy (Figure 3D) and the degree of white fat deposition into the BAT (Figure 3E). Although not statistically significant, the average size of pancreatic Langerhans islets was slightly higher in *Cdc14a* KO mice when compared to wild type (Figure 3F). Overall, these results suggest that elimination of CDC14A causes an exacerbation of the physiological response of mice to HFD treatment and suggest differential effect of the two CDC14 phosphatases in the control of adipogenesis.

**Figure 3.**
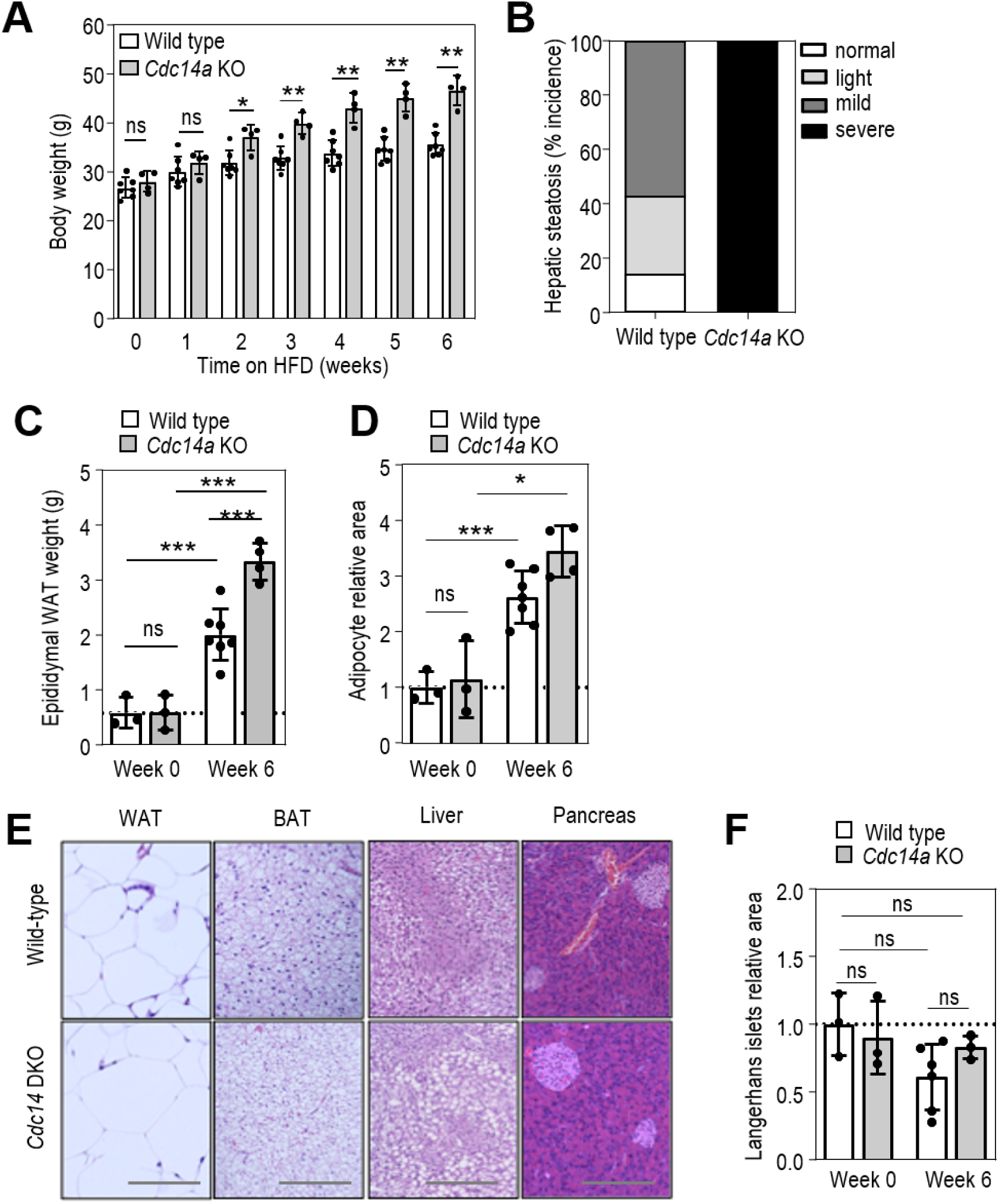
Elimination of CDC14A exacerbates high fat diet-induced obesity. **A**. Body weight of wild type and Cdc14a-null mice after 6 weeks of high fat diet (HFD). **B**. Incidence of hepatic steatosis in wild type and Cdc14a KO mice at the end of the HFD treatment. **C**. Weight of the epidydimal WAT of wild type and Cdc14a KO mice after HFD treatment. Each point represents the average relative size area of at least 50 adipocytes of each mouse. **D**. Quantification of the average area of the adipocyte cells that composed the WAT in wild type and *Cdc14*a KO mice after HFD. Each point represents the average relative size area of at least 50 adipocytes of each mouse. **E**. Representative H&E staining of WAT, BAT, liver and pancreatic tissue sections of wild type and Cdc14a KO mice at the end of HFD treatment. N=7 wild type, N=4 Cdc14a KO for HFD treatment; N=3 wild type, N=3 Cdc14a KO mice for control mice in normal diet. All mice used in this experimental setting were male between 15 and 18 weeks of age. Scale bars, 100 μm. **F**. Quantification of pancreatic B-islets in wild type and Cdc14a KO mice after 6 weeks of HFD or in control animals never subjected to HFD. Each dot represents the average relative area of at least 7 B-islets per mice. In A,C,D,F, data are shown as mean ± SEM. Statistical significance was assessed using 2-way ANOVA (Bonferroni’s multiple test) for A, B and C, and the two-tailed unpaired Student’s t-test with Welch correction F. ns, not significant (p>0.05); *, p<0.05; **, p<0.005; ***, p<0.001, ****, p<0.0001.

### Cdc14 DKO mice are resistant to high fat diet-induced obesity and glucose intolerance

We next submitted *Cdc14* DKO adult mice to HFD for six consecutive weeks. As expected, control animals showed a significant increase in their body weight. Unexpectedly, *Cdc14* DKO mice weight nearly did not change under this treatment (Figure 4A). After six weeks of treatment, control animals presented a cumulative increase of 30% in their body weight, while *Cdc14* DKO animals increased only by 10% (Supplementary Figure 2A,B). Concomitantly, accumulation of fat in the form of WAT (epidydimal region) was four times lower in *Cdc14* DKO animals when compared with wild type ones (Figure 4B). As expected, histological analysis showed hepatic steatosis, adipocyte hypertrophy and white fat deposition into brown adipose tissue in the control group (Figure 4C,D,E). In contrast, *Cdc14* DKO mice revealed decreased incidence and severity of hepatic steatosis (Figure 4C), diminished white fat deposition into brown adipose tissue (Figure 4E) and adipocyte cells display an area size approximately 3 times smaller than wild type adipocytes at the end of the HFD treatment (Figure 4D). The expected reduction in pancreatic Langerhans islets after the HFD treatment was lower in knockout than in wild type mice, although statistical significance was not reached (Supplementary Figure 2C). In view of these results, glucose (GTT) and insulin (ITT) tolerance tests were performed and showed that *Cdc14* DKO mice were protected against HFD-induced glucose intolerance (Figure 4F and Supplementary Figure 2D).

**Figure 4.**
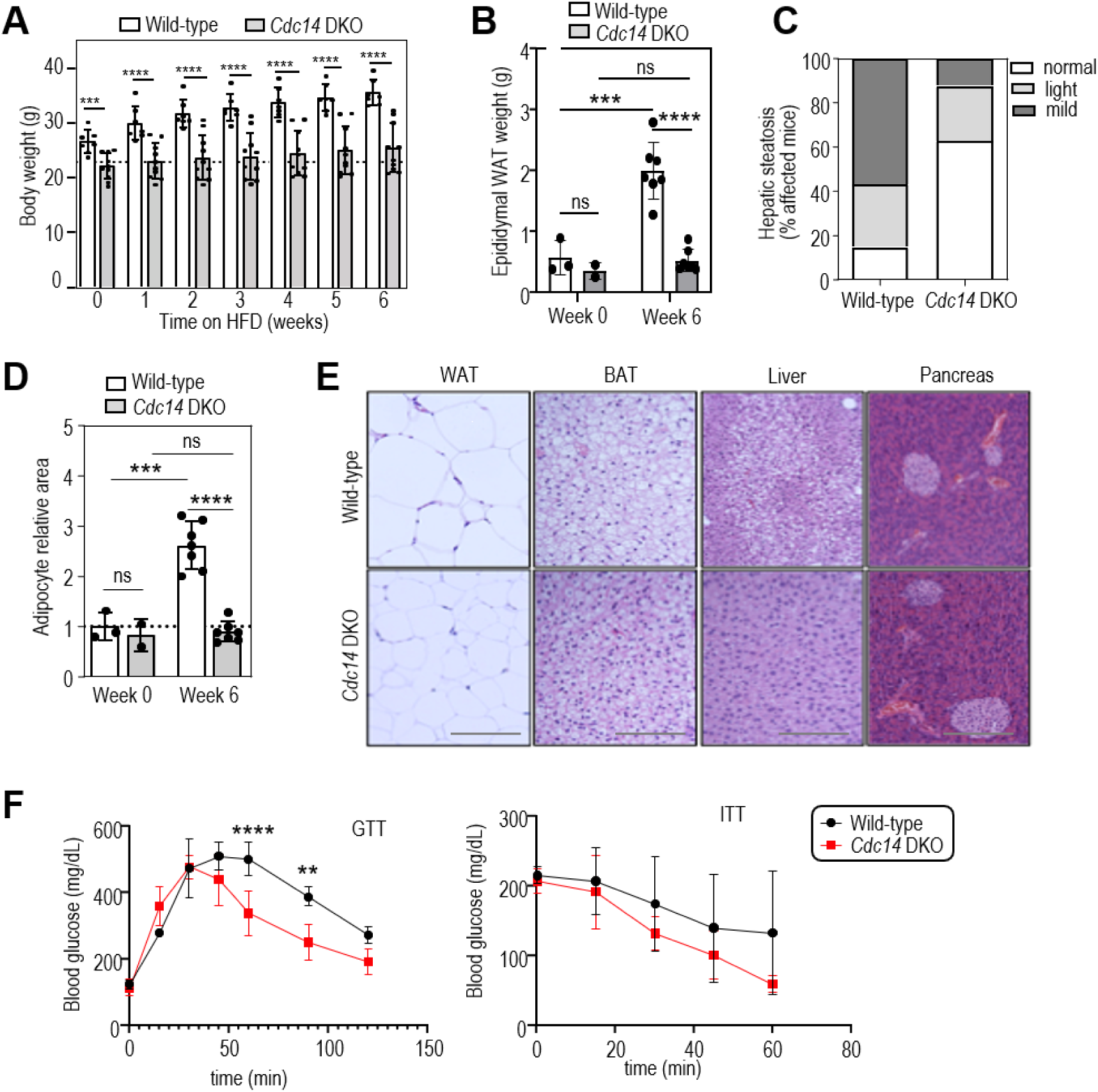
*Cdc14* DKO mice are resistant to high fat diet-induced obesity. **A**. Body weight of wild type and Cdc14 DKO mice after 6 weeks of high fat diet (HFD). **B**. Weight of the white adipose tissue (WAT) in the gonadal area of wild type and Cdc14 DKO mice after the HFD treatment. p<0.0001. **C**. Incidence of hepatic steatosis in wild type and *Cdc14* DKO mice at the end of the HFD treatment. **D**. Quantification of the average area of the adipocyte cells that composed the WAT in wild type and *Cdc14* DKO mice after HFD. Each point represents the average relative size area of at least 50 adipocytes of each mouse. **E**. Representative H&E staining of WAT, BAT, liver and pancreatic tissue sections of wild type and *Cdc14* DKO mice at the end of HFD treatment. N=7 wild type, N=7 *Cdc14* DKO for HFD treatment; N=3 wild type, N=3 *Cdc14* DKO mice for control mice in normal diet. It should be noted that from an initial experimental group of 10 mice, 3 Cdc14 DKO mice behaved like wild type and gained weight normally. These mice were regarded outliers and were not considered for the subsequent analysis abovementioned. All mice used in this experimental setting were male between 15 and 18 weeks of age. Scale bars, 100 μm. Data are shown as mean ±SEM. Statistical significance was assessed using 2-way ANOVA (Bonferroni’s multiple test) for A and the two-tailed unpaired Student’s t-test with Welch correction for B-D (p>0.05); *, p<0.05; **, p<0.005; ***, p<0.001, ****. **F**. Glucose tolerance test (GTT) and insulin tolerance test (ITT) of wild type and *Cdc14* DKO mice at the end of the HFD treatment. N=4 for wild type, N=5 *Cdc14* DKO.

Since changes in body weight and fat mass can be caused by variations in energy expenditure due to decreased nutrient ingestion or increased activity related with abnormal motion patterns, wild-type and *Cdc14* DKO mice were subjected to a comprehensive animal metabolic monitoring system. Oxygen consumption (VO2), carbon dioxide production (VCO2), energy expenditure (EE) and respiratory quotient (RQ) were measured throughout 72 hours during the dark and light periods and did not differ significantly between wild type control and *Cdc14* DKO mice (Supplementary Figure 3). The spontaneous activity of the mouse along the cage showed no major differences between the two groups, although *Cdc14* DKO mice exhibited increased rearing activity during the night period, as well as increased water intake (Supplementary Figure 3).

### CDC14 phosphatases regulate adipogenic differentiation

To investigate the cellular and molecular basis of the effect of CDC14 phosphatases, we induced adipocyte differentiation *in vitro*^22^ in wild-type mouse embryonic stem cells (ESCs) and in ESCs depleted of CDC14A and B (individually or in combination) using CRISPR-Cas9 technologies. Cells individually depleted of CDC14A or CDC14B showed a strong reduction in the mRNA levels of the corresponding transcript (Figure 5A). Worthy of note, *Cdc14a* knockout resulted in upregulation of *Cdc14b* transcripts and vice versa (Figure 5A), suggesting compensatory mechanisms. 24 days after adipogenesis induction lipid content was measured by Oil Red staining, showing that single depletion of CDC14B produced a drastic reduction in differentiation of almost 70%, and this effect was not further increased after co-depletion of both CDC14A and CDC14B (Figure 5B). In accordance with the results obtained *in vivo,* the ability of ESCs to differentiate into adipocytes was significantly enhanced by approximately 45% upon single depletion of CDC14A. This effect was very likely due to the overexpression of CDC14B as it was rescued by genetic ablation of *Cdc14b* (Figure 5B).

**Figure 5.**
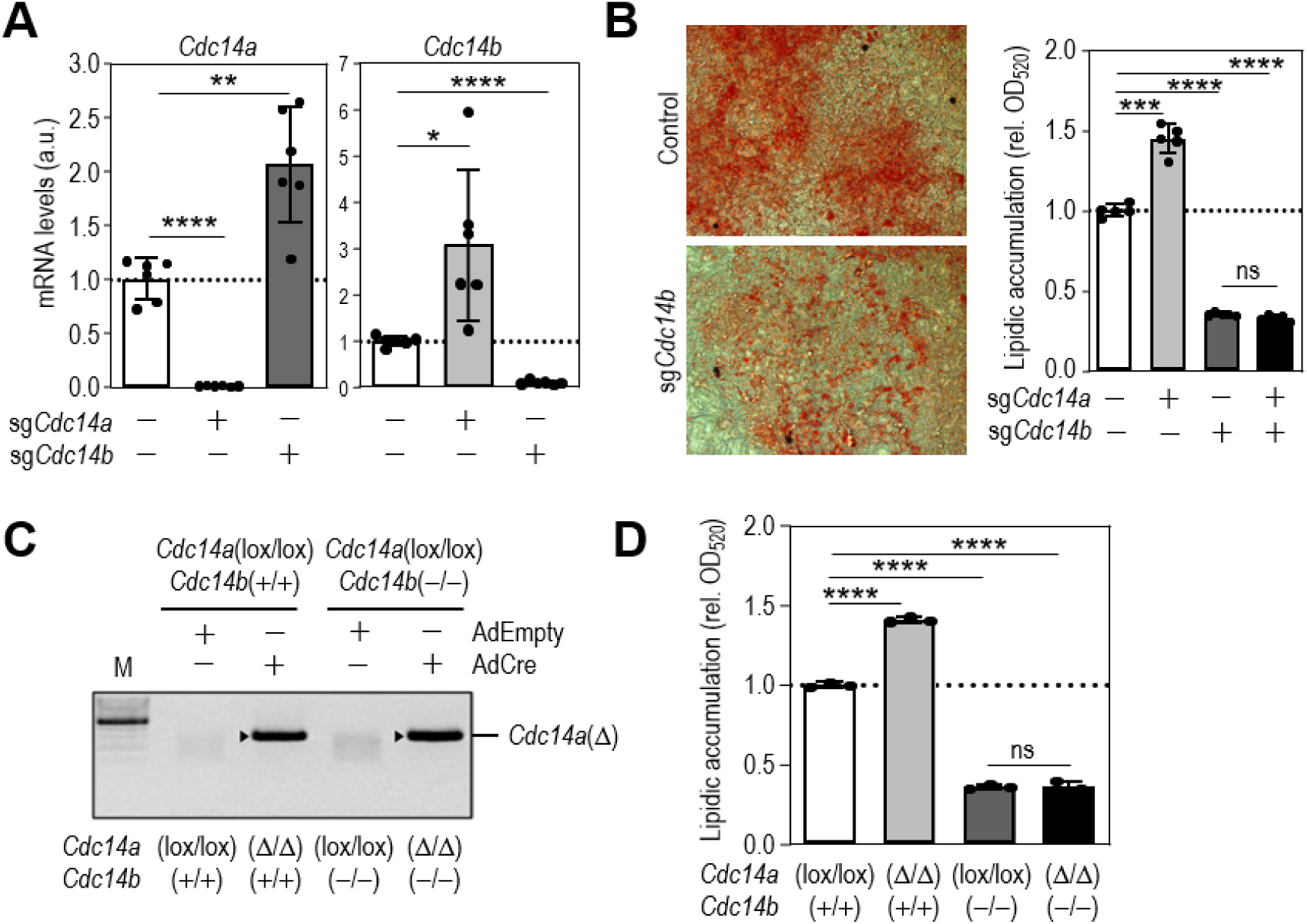
CDC14 phosphatases regulate adipogenic differentiation. **A**. mRNA expression analysis of *Cdc14a* and *Cdc14b* levels in mESCs wild type or depleted of CDC14A or CDC14B by CRISPR-Cas9 technology. **B**. Representative images and quantification of lipid accumulation by Oil Red O staining of wild type ESCs or ESCs depleted of CDC14a, CDC14b or CDCA+B by CRISPR/Cas9 at day 24 of the adipocyte differentiation protocol. **C**, Representative amplification of PCR products of the Cdc14a-null locus of mESCs infected with adenoviruses expressing Cre recombinase (AdCre) or, as a control, empty adenoviruses (AdEmpty). **D**, Oil red O quantification of lipid accumulation in mESCs control [*Cdc14a*(lox/lox);*Cdc14b*(+/+)], *Cdc14a* KO [*Cdc14a*(Δ/Δ);*Cdc14b*(+/+)], *Cdc14b KO* [*Cdc14a* (lox/lox);*Cdc14b*(Δ/Δ)] and *Cdc14a+b* KO [*Cdc14a*(Δ/Δ);*Cdc14b*(Δ/Δ)) after 24 days of the adipocyte differentiation protocol. N=3 independent experiments. Data are means ± SD. Statistical significance was assessed using the two-tailed unpaired Student’s t-test with Welch correction. ns, not significant (p>0.05), *, p<0.05; **, p<0.01; ***, p<0.001, ****, p<0.0001.

To further confirm these results and to address the discrepancy between the *in vivo* and *in vitro* data concerning the single elimination of CDC14B, we performed *in vitro* differentiation experiments with ESCs directly obtained from the genetically-modified mouse models. To avoid potential long-term compensatory effects, we used a conditional *Cdc14a* knockout allele [*Cdc14a*(lox/lox) ESCs cells)] in combination with the constitutive *Cdc14b* KO [*Cdc14b*(−/−)] model. Acute genetic deletion of *Cdc14a* [*Cdc14a*(Δ) allele] was achieved after infection of these cells with adenoviruses expressing Cre recombinase (AdCre), thus generating the four genotypes of our interest: wild-type [*Cdc14a*(lox/lox); *Cdc14b*(+/+)], conditional *Cdc14a* ablation [*Cdc14a*(Δ/Δ); *Cdc14b*(+/+)], *Cdc14b* KO [*Cdc14a*(lox/lox); *Cdc14b*(−/−)] and conditional *Cdc14a* ablation in a *Cdc14b*-null background [*Cdc14a*(Δ/Δ); *Cdc14b*(−/−)] (Figure 5C). These experiments confirmed that depletion of CDC14A enhances adipogenic potential, while *Cdc14b* KO cells have diminished adipocyte differentiation ability, an effect that is not altered by concomitant ablation of *Cdc14a* (Figure 5D).

We have previously described that CDC14B modulates neural differentiation, in part through the de-phosphorylation of the undifferentiated transcription factor 1 (UTF1)^12^. To eliminate the possible effect of UTF1 in adipocyte differentiation from ESCs, we also used a model of induced adipocyte trans-differentiation from primary mouse embryonic fibroblasts (MEFs) as UTF1 is not expressed in these differentiated cells. Ten days after exposure to inducers, Oil Red O staining demonstrated enhanced differentiation of *Cdc14a* KO MEFs with an increase degree of adipose conversion of approximately 42% compared to wild type MEFs (Supplementary Figure 4). Both *Cdc14b* KO and *Cdc14* DKO MEFs exhibited a reduction in differentiation, of 21% and 33%, respectively, although the differences were not statistically significant. Remarkably, adipogenic differentiating assays using primary MEFs have very low efficiency (15–30% of adipocytes), and this variable heterogeneity might generate background noise and partially mask the effect of CDC14 depletion (Supplementary Figure 4). These experiments suggest that CDC14 phosphatases affect adipocyte differentiation by mechanisms that are independent of UTF1.

### CDC14B dephosphorylates PPARγ allowing adipocyte differentiation

To investigate the mechanisms by which CDC14 phosphatases might be controlling adipocyte differentiation, we performed transcriptomics analysis at days 0, 5 and 35 of the differentiation protocol using wild-type ESCs or ESCs depleted of CDC14A and CDC14B (individually or in combination) (Figure 6A and Supplementary Figure 5A). Gene expression analysis showed the upregulation of proteins involved in adipocyte differentiation with time, and the concomitant downregulation of cell cycle proteins as well as stem-cell markers (Figure 6B and Supplementary Figure 5B). *Cdc14b* KO and *Cdc14* DKO cells, however, displayed deficient accumulation of genes involved in adipocyte differentiation and maintained a more active proliferation state when compared to control cells Figure 6B,C and Supplementary Figure 5C). *In silico* analysis of the main transcription factor controlling deregulated genes in *Cdc14b* KO cells pointed to the stress-response transcription factors TP53, as well as SUZ12 (Suppressor of Zeste 12 Homolog), a core component of the Polycomb Repressive Complex 2 (PRC2), that plays a crucial role in stem cell differentiation by repressing gene expression^23^. In addition to these genes, PPARγ, a key protein in the adipogenesis process, was enriched in the promoters of genes deregulated in *Cdc14b*-null cells and the PPARγ signaling pathway display a reduced activation in *Cdc14b*-deficient cells (Figure 6D).

**Figure 6.**
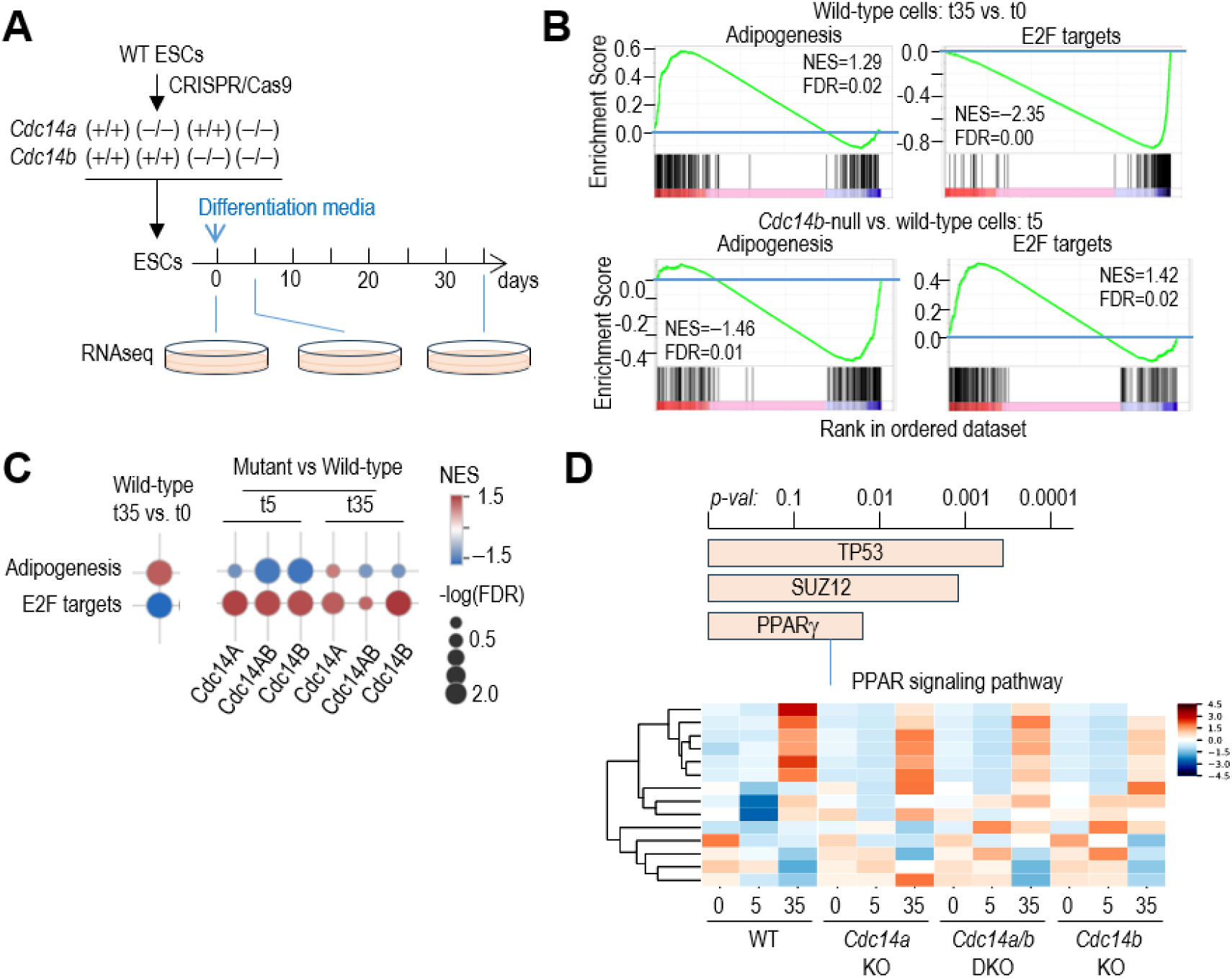
CDC14B is required for proper regulation of genes involved in differentiation and the PPARγ signaling pathway. **A,** Schematic representation of the experimental procedure carried out for the RNAseq experiment with ESCs wild-type or depleted of CDC14A and/or B during adipogenic differentiation. **B**, GSEA plots of the Hallmarks genesets involved in Differentiation or cell cycle (E2F targets) in the conditions indicated. **C**, Normalized enrichment score (NES) and false discovery rate (FDR) for the indicated comparisons between CDC14-mutant and control cells at different time points during the differentiation process. Changes between 35 and 0 days in the control cells are shown to the left as a control. **D**, Analysis of enrichment of regulatory sequences (based on ENCODE and ChEA transcription factor signals) in genes deregulated in Cdc14b-null cells after differentiation. The bottom panel shows a heat-map analysis of genes involved in the PPARγ signaling pathway obtained by RNA-seq of the above-mentioned samples.

PPARγ has been previously shown to be phosphorylated at several residues, with different effects for the functionality of the protein^24^. Therefore, we analyzed whether CDC14B could directly interact with and dephosphorylate PPARγ by using co-immunoprecipitation and coupled kinase-phosphatase *in vitro* assays. PPARγ1 and CDC14B co-immunoprecipitate in the chromatin-bound fraction, suggesting that both proteins are able to interact (Supplementary Figure 6A). As expected, ERK2 was able to phosphorylate PPARγ, and incubation of phosphorylated PPARγ with CDC14B −but not with a phosphatase-dead version (CDC14B-PD) or with CDC14A− decreased PPARγ phosphorylation status (Figure 7A). In addition, western blot analysis of these *in vitro* kinase-phosphatase assays using cold ATP, revealed that CDC14B is able to dephosphorylate PPARγ at S/TP motifs (Figure 7B). Furthermore, phospho-residue specific antibodies indicated that CDC14 dephosphorylates serine 112 but not serine 273 (Figure 7C). To confirm this, we subjected PPARγ phosphorylated by ERK2 or CDK5 and treated with CDC14B or the phosphatase-dead (CDC14-PD) isoform to phospho-proteomic analysis. As expected, we observed that S112 phosphorylation induced by ERK2 is decreased by CDC14B but not by CDC14B-PD (Figure 7C). Surprisingly, the phosphorylation of S273, induced either by ERK2 or CDK5, also decreased (Figure 7C). In addition, we observed that T296 was exclusively phosphorylated by CDK5 but not ERK2, and this residue was also de-phosphorylated by CDC14B. Finally, T75 was only phosphorylated by ERK2 and not CDK5, and this phosphorylation was not affected by CDC14B (Supplementary Figure 6B).

**Figure 7.**
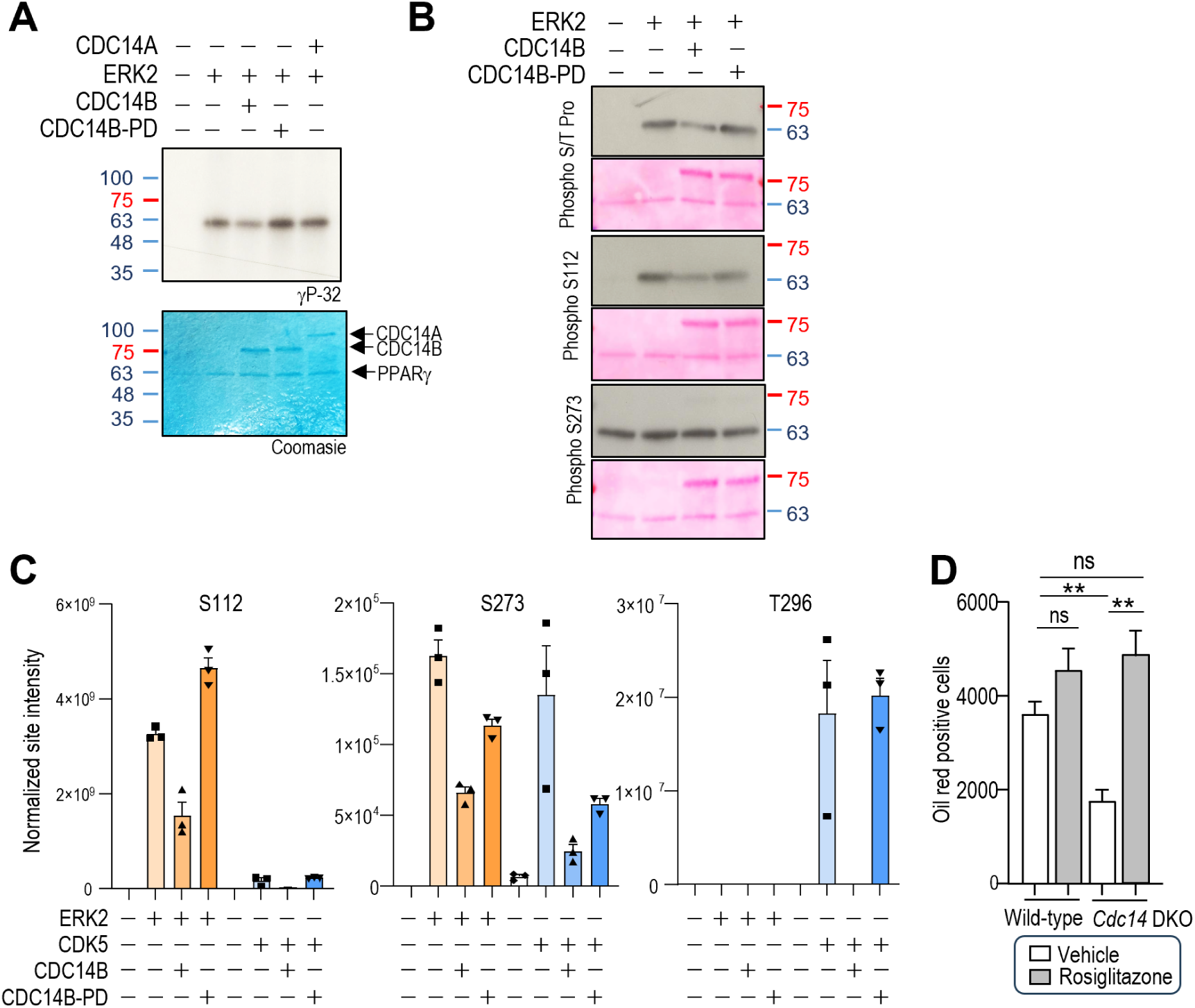
CDC14B dephosphorylates PPARγ allowing adipocyte differentiation. **A**. *In vitro* phosphatase assay of PPARγ2 previously phosphorylated by ERK2 and subsequently exposed to wild-type or a catalytically inactive form (PD) of CDC14A or B. **B**, Western blot analysis of PPARγ2 after *in vitro* kinase phosphorylation. **C**. Phospho-proteomic analysis of ERK2 or CDK5-induced phospho-PPARγ2 at residues S112, S273 and T296 after incubation with CDC14B or CDC14B-PD. **D**. Quantification of lipid accumulation by Oil Red O staining in wild type and *Cdc14a+b* KO mESCs after 24 days of adipogenic differentiation in presence or absence of troglitazone. N=3 independent experiments. Graph are means ± SD. Statistical significance was assessed using the two-tailed unpaired Student’s t-test. **p<0.005.

It has been previously described that S112 phosphorylation inhibits PPARγ adipogenic activity^25^, which might explain why adipogenic differentiation is impaired in CDC14 DKO cells. To test whether PPARγ mediates the effects of CDC14 deficiency, we increased PPARγ activity in *Cdc14* DKO differentiating cells by treating them with troglitazone, a PPARγ agonist that has also been reported to block its phosphorylation in S273^21^. Interestingly, the increase in PPARγ activity by troglitazone was able to compensate for the lack of CDC14 phosphatases and rescued adipogenic differentiation in *Cdc14* DKO cells (Figure 7D), suggesting that CDC14 controls adipogenic differentiation by regulating PPARγ activity.

## Discussion

The role of CDC14 phosphatases in vertebrates has long been incompletely understood, partly due to the functional redundancy between isoforms^26–28^. Some of the functions described for these phosphatases include regulation of centrosome and spindle function, ciliogenesis, transcription and DNA damage^5–11,27–29^. We recently generated a mouse model with simultaneous depletion of CDC14A and CDC14B. Primary cells derived from these mice did not show defects in mitosis exit or proliferation. However, DKO mice exhibited partial embryonic lethality and perinatal mortality accompanied with defects in neurogenesis, suggesting a new role of CDC14 phosphatases in the differentiation of stem cell into neural lineages^12^. This effect is at least partially mediated by the modification of UTF1, an epigenetic regulator exclusively expressed in undifferentiated cells. During differentiation, CDC14 dephosphorylates UTF1 leading to its ubiquitin-dependent destabilization and the subsequent expression of genes with bivalent promoters^12^.

Intriguingly, CDC14 DKO mice that survive the perinatal period do not show major defects in lifespan. However, our results suggest that these mutant mice do not gain weight with age and are resistant to HFD-induced obesity and glucose intolerance, also exhibiting decreased hepatic steatosis. Of note, some *Cdc14* DKO mice can also escape this phenotype, confirming the variable penetrance of CDC14 depletion, possibly by compensating phosphorylation levels through alternative pathways. *In vitro* studies with ESCs and MEFs indicate that concomitant depletion of CDC14A and CDC14B decreases adipogenesis in a UTF1-independent manner. Transcriptomic and biochemical studies suggest that CDC14B directly de-phosphorylates PPARγ, a master regulator of adipogenesis, and the PPARγ agonist troglitazone can rescue adipogenesis *in vitro* in *Cdc14* DKO cells. PPARγ is also important for neurogenesis, regulating neural stem cell proliferation and differentiation^30^, and PPARγ agonists like thiazolidinediones do not only increase adipogenesis and insulin sensitivity, but also reduce neurodegeneration in mouse models of encephalomyelitis and Alzheimer disease^31,32^, although its final effect in improving cognitive function is controversial^33^. Therefore, a possible contribution of PPARγ in the neural defects observed in the *Cdc14* DKO mice cannot be excluded.

We have observed in this work differential effects when analyzing the individual contribution of CDC14A and CDC14B. Depletion of both enzymes impairs adipogenesis *in vivo* and *in vitro*, whereas individual depletion of CDC14B impairs adipogenesis *in vitro* but not *in vivo*, suggesting that a compensatory mechanism can be activated at the organismal level. Intriguingly, CDC14A depletion increases both adipogenesis *in vitro* and *in vivo*, resulting in increased weight under HFD. This effect might be explained by the fact that CDC14B is upregulated in *Cdc14a*-null cells, possibly leading to decreased levels of PPARγ phosphorylation and, consequently, increased activity.

PPARγ function is tightly regulated by posttranslational modifications, including phosphorylation at residues S112, S273, S133, T75 and T296 with different consequences for PPARγ activity. As stated above, S112 can be phosphorylated by ERK kinases, resulting in a reduced activity by altering its ability to recruit different co-repressors or co-activators without disturbing its stability or DNA-binding ability^20^. However, the consequences of S112 phosphorylation seem opposite when mediated by CDK7 or CDK9, suggesting context-dependent effects^24^. Importantly, the S112A allele correlates with a decreased risk of type 2 diabetes^34^. Here we observe that S112 can be directly de-phosphorylated by CDC14B, which was also previously described to be de-phosphorylated by PP5, PPM1B and WIP1^35–37^, although the relevance of these other phosphatases in adipogenesis or glucose sensitivity *in vivo* has not been shown so far.

PPARγ S273 is phosphorylated by CDK5 leading to decreased expression of a subset of PPARγ target genes related to insulin sensitivity without affecting PPARγ adipogenic activity^38^. Increased S273 phosphorylation is observed in obesity and can be counteracted by PPARγ agonists, thereby improving metabolic profiles of both mice on HFD and patients with impaired glucose tolerance. Interestingly, PPARγ binding compounds that decrease S273 phosphorylation with no or little agonistic effect on PPARγ, also have similar metabolic effects with less side effects than full agonists such as TZDs^39^. Thus, blocking S273 phosphorylation has interesting clinical applications. Here we show that CDC14B can directly dephosphorylate S273, identifying for the first time a phosphatase of this critical residue. We have also detected that the T296 residue, exclusively phosphorylated by CDK5 but not ERK2, can be dephosphorylated by CDC14B, revealing that it is the main CDK5-counteracting phosphatase in PPARγ identified so far. However, the exact contribution of each residue to the final phenotype remains unknown.

In summary, we have demonstrated that CDC14 phosphatases regulate adipogenesis and glucose intolerance *in vivo*. Given the pivotal role of adipogenesis in metabolic regulation and the central function of PPARγ in this process, our findings could have significant clinical implications for metabolic disorders such as type 2 diabetes. Furthermore, this study reinforces the hypothesis that cell cycle regulators may have evolved in multicellular organisms to mediate the intricate cross-talk between proliferation and differentiation, thereby supporting the complexity required for tissue specialization and organismal development.

## STAR Methods

### Mouse models and high-fat diet

The *Cdc14a* and *Cdc14b* double knockout (DKO) mouse model has been described previously in Villarroya-Beltri et al^12^. Work with these mice were approved by the corresponding ethical committees at the Instituto de Salud Carlos III and Comunidad de Madrid.

For the High Fat Diet (HFD) experiments performed in this study, mice were fed for 6 weeks with a 45% high fat diet (45% fat, 20% protein, 35% carbohydrate; Research Diets; D12451). All mice used for such treatment were males with an age comprised between 15 and 18 weeks. Mouse body weight was assessed weekly. At the end of experiment (6 weeks), both body weight and epididymal white adipose tissue (WAT) weight were measured and tissue samples of liver, pancreas, WAT and brown adipose tissue (BAT) were acquired for detailed histological assessment in both the control group and mutant mice. Additionally, control samples of mice that have not been fed with HFD (week 0) were also acquired for comparison.

### Metabolic assays

The Oxylet (Harvard Apparatus) was used to measure: indirect calorimetry, food/water intake and locomotive activity. Indirect calorimetry is a method used to measure the energy consumption of animals. The respiratory quotient (RQ) was calculated as RQ=CO2 production/ Volume O2 consumption. Energy expenditure (EE) was calculated by taking advantage of the semi-empirical Weir Equation. Locomotive activity was measured by analyzing mean spontaneous activity of the mouse along the cage and rearing activity. All mice used in this experiment were males with an age comprised between 17 and 20 weeks. Animal experimentation occurred over a period of 3 days. All data presented is normalized relatively to the total body weight of each mouse.

For Insulin tolerance test (ITT) mice were fasted for 6h and then intraperitoneally injected with rapid-acting insulin (0.75 U/kg body weight) (Actrapid, Novo Nordisk), followed by tail vein blood glucose sampling to measure glycemia at 0, 15, 30, 45, and 60 min after insulin injection. Glycemia was measured in blood samples obtained from lateral tail vein with a glucometer and glucose strips (Accu-Chek Instant, Roche). Glucose tolerance test assay (GTT) was performed in mice fasted for 16 h and then administered an intraperitoneal glucose injection (2 g/kg body weight).

### Histological processing

For histological analysis, tissue samples were fixed overnight in 10% buffered formalin (Sigma-Aldrich). Samples were paraffin-embedded and cut into 3-5μm thickness sections that were mounted in superfrost®plus slides and dried overnight. Sections were then stained with hematoxylin and eosin (H&E). Image acquisition was performed using a Leica D3000 microscope. For adipocyte size quantification a minimum of 50 cells per mice were quantified in 20X field image pictures. Size of pancreatic B-islets is the average of at least 8 different islets per mice.

### Cell culture

Mouse Embryonic Stem Cells (ESCs) were cultured over mitomycin C (Roche)-inactivated mouse embryonic fibroblasts (MEFs) and in the presence of mESC medium containing KO-DMEM (Gibco), 2-Mercaptoethanol (Invitrogen), non-essential aminoacids MEM NEAA (Invitrogen), GlutamMAX (Gibco), Penicillin/Streptomycin (5000 μg/ml, Invitrogen), LIF (Leukemia Inhibitor Factor, ESGRO, Millipore) and 10% Fetal Calf Serum (Hyclone) at 37°C in a humidified 5% CO2 atmosphere. Cell medium was changed every 24h.

For lentiviral transduction HEK-293T cells were transfected using Lipofectamine 2000 (Invitrogen) with the lentiviral vector of interest along with packaging pMDL, REV and envelope (VSVG) expression plasmids according manufacturer’s instructions. Virus-containing supernatants were collected at 48 and 72h after transfection and used to transduce cells of interest in the presence of polybrene (4 μg/mL) in two consecutive rounds of infection. For lentiviral transduction of ESCs, viral supernatants were produced in mESCs medium, and mESCs were seeded in 0.1% gelatin-coated plates 48h before transduction. 36h after transduction, cells were selected using puromycin (0.5 μg/mL) during 2 days and subsequently transferred back to culture over feeders. For adenoviral transduction, adenoviruses expressing CMV-Empty or CMV-Cre (Cre recombinase) were obtained from The University of Iowa (Iowa City, IA) and infection was carried out at a 250 MOI for two consecutive rounds 12 hours apart in 6-well plates coated with 0.1% gelatin.

### Gene editing by CRISPR/Cas9

Single guide RNAs (sgRNAs) targeting exon 4 of the Cdc14a allele and exon 8 of the *Cdc14b* allele were designed according to available algorithms (http://crispr.mit.edu/) and cloned into the BmsbI site of the lentiCRISPR V2.0 backbone vector (Addgene, plasmids# 52961). Lentiviral vector containing the corresponding sgRNAS or the lentiCRISPR empty vector as control were used to transduce cells of interest as described above. Cutting efficiency of the guides was assessed by the T7E1 assay according with the method described by Zhang laboratories (https://zlab.bio) and using specific primers (Supplementary Table 1).

### Adipogenic differentiation in vitro

Primary MEFs were differentiated into adipocytes upon exposure to hormonal inducers as reported by Zhang et al^40^. Briefly, primary MEFs at passage 2 were split in 6-well plates and cultured until reaching confluence. Two days later, cells were treated with adipogenic differentiation induction medium containing DMEM, 0.5 mM methylisobutylxanthine (IBMX), 1 uM dexamethasone (DEX), 10 μg/ml insulin, 10 μM troglitazone, 10% (vol/vol) FBS and gentamicin. Two days after, the medium was replaced by maintenance medium containing DMEM, 10 μg/ml insulin, 10 μM troglitazone, 10% (vol/vol) FBS and gentamycin. The medium was renewed every other day. Adipocytes colony formation and lipidic accumulation were evaluated by oil red staining 10 days after differentiation induction.

Differentiation of ESCs to mature adipocytes was performed using adipogenic inducing cocktails as described by Cuaranta-Monroy et al. with modifications^22^. Cells were dissociated using Trypsin to achieve a single cell suspension and resuspended to a concentration of 2×10^5^ cells/ml in the presence of complete growth medium lacking leukemia inhibitory factor (LIF). Small drops of this suspension (∼30 μl) were collected and seeded on the lid of a 10 mm plate that was then inverted and placed over the bottom of a bacteriological Petri dish filled with 10 mL of PBS, generating hanging drops. Cells were incubated at 37°C in a humidified 5% CO2 incubator. After 4 days, embryoid bodies (EB) were visible at the bottom of the drops and the EB suspension was transferred to a non-adherent gelatin-coated 6 well culture plate containing ES differentiation medium. Two days later, cells were treated with RA (1 μM) and AsA (12.5 μg/mL) for 3 days and in the fourth day only RA (1μM) was used. Cells were then treated for a week with an adipogenic cocktail containing: Insulin (0.5 μg/mL), T3 (3 nM) and AsA (12.5μg/mL). Medium was changed daily. Cells were treated with a second adipogenic cocktail for an extra week containing: IBMX (0.5 mM), Dexamethasone (0.1 μM), Insulin (20 μg/mL), Indomethacin (0.06 mM) and AsA (25 μg/mL). When indicated, troglitazone was added to the differentiation media. Medium was changed every two days. Finally, medium was supplemented with Insulin (10 μg/mL), AsA (25 μg/mL) and T3 (3 nM) for six additional days, time point at which (day 30) cell cultures were evaluated for adipocyte colonies and lipidic accumulation by oil red O staining.

### Oil red staining

Cells were washed twice carefully with PBS and fixed with 4% paraformaldehyde for 1 hour at RT. Cells were then washed twice with distilled water and incubated for 5 minutes with 60% isopropanol. For Oil red O staining solution preparation, 3 parts of Oil red O dye (Sigma Aldrich) were mixed with 2 parts of distilled water and allow to sit at RT for 15 minutes. The solution was then filtered immediately used to cells by incubating for 1 hour at RT. Excess staining was removed by washing twice with distilled water. Image acquisition was performed before and after staining, using a Leica D3000 microscope. Image analysis was done using ImageJ software. Spectrophotometric quantification of the stain was performed by dissolving the stained oil droplets in 100% isopropanol for 10 minutes. Optical density was then measured at 520 nm.

### RNA expression analysis

To quantify RNA expression by qPCR, total RNA from cells and tissues was isolated using Trizol (Invitrogen) according to the manufacturer’s instructions. Reverse transcription from 1 μg of total RNA was performed with the M-MLV retrotranscriptase (Promega) followed by quantitative PCR using SYBR Green PCR Master Mix (Applied Biosystems). Oligonucleotides used are listed in Supplementary Table 1.

For RNAseq, total RNA was extracted using miRVANA RNA extraction kit (Ambion). 500 ng of total RNA was used. RNA Integrity Number was analyzed on Agilent 2100 Bioanalyzer (range 9.5–10). Sequencing libraries were prepared with the “QuantSeq 30 mRNA-Seq Library Prep Kit (FWD) for Illumina” (Lexogen, 015) following manufacturer instructions. Briefly, library generation is initiated by reverse transcription with oligodT priming, followed by a random-primed second strand synthesis. Primers from both steps contain Illumina-compatible sequences. The libraries were completed by PCR and sequenced on Illumina HiSeq2500. Sequencing quality control was performed using fastqc preprocessing (v0.11.9) and adapters were removed with BBDuk (bbtools v37.62). For each sample, we performed read alignment to the Gencode version of GRCh38 mouse reference genome using STAR (v2.7.10a), indexing using samtools (v1.14) and quantification using htseq-count (v1.99.2) and finally, count matrices were extracted for downstream analysis.

The resulting reads were analyzed with the nextpresso pipeline^41^ or routine pipelines in R and python. Raw count data were normalized using the median ratio normalization method, and lowly expressed genes were filtered out before analysis. Differential gene expression analysis between condition groups was performed using the DESeq2 R Bioconductor package (v1.32.0)^42^. Gene expression differences were assessed using the Wald test, generating log2 fold change (LFC) values, standard errors, p-values, and adjusted p-values using the Benjamini-Hochberg correction method to control the false discovery rate (FDR). Genes with an adjusted p-value (padj) < 0.05 were considered significantly differentially expressed. GSEA Preranked was used to perform gene set enrichment analysis for the selected gene signatures on a pre-ranked gene list, setting 1,000 gene set permutations^43^ using the Python wrap^44^. Gene sets with significant enrichment levels (FDR < 0.25) were considered.

### Biochemical studies

HeLa cells were transfected with CDC14B and PPARγ using Lipofectamine 2000 (Invitrogen). Cultured cells were harvested and lysed in RIPA (25 mM Tris–HCl pH 7.5, 150 mM NaCl, 1% NP-40, 0.5% Na deoxycholate, Complete protease inhibitor cocktail) 15 min at 4°C and centrifuged 10 min at 14,000 g. Supernatant containing cytoplasmic and nuclear-soluble proteins was saved, and pellet containing chromatin-bound and nucleolar proteins was resuspended in high-salt RIPA (25 mM Tris–HCl pH 7.5, 500 mM NaCl, 1% NP-40, 0.5% Na deoxycholate, complete protease inhibitor cocktail) incubated for 10 min at 4°C and centrifuged 10 min at 14,000 g. Protein G Dynabeads (Thermo fisher) were washed in PBS-tween 0.01%, incubated with anti-GFP antibody for 1 h at RT, and equilibrated in RIPA buffer. RIPA or high-salt RIPA protein extracts were incubated with pre-washed and equilibrated Dynabeads 1 h at 4°C for pre-clearing. Pre-cleared protein extracts were incubated with antibody-coupled Dynabeads for 2 h at 4°C. Dynabeads were washed five times with RIPA buffer, resuspended in loading buffer, and boiled for 5 min. Dynabeads were finally removed, and protein-containing extracts were loaded for Western blot analysis.

For immunoblot analysis, protein lysates were mixed with loading buffer (350 mM Tris–HCl pH 6.8, 30% glycerol, 10% SDS, 0.6 M DTT, 0.1% bromophenol blue) boiled for 5 min, and subjected to electrophoresis using the standard SDS–PAGE method. Proteins were then transferred to a nitrocellulose membrane (BioRad), blocked for 1h at RT in TBS 0.1% Tween-20 containing 5% BSA, and incubated overnight at 4°C with specific primary antibodies. Membranes were washed 10 min in TBS-Tween and incubated with peroxidase-conjugated secondary antibodies (Dako, 1:5,000) for 45 min at RT. Finally, the membranes were washed for 5 min three times and developed using enhanced chemiluminiscence reagent (Western Lightning Plus-ECL; Perkin Elmer). The following primary antibodies were used: GFP (Roche, 11814460001), Phospho-Ser273 PPARγ (Thermo Scientific, BS-4888R), Phospho-Ser112 PPARγ (Thermo Scientific, PA5-36763), PPARγ (Cell Signaling Technology 2435S), Phospho-(Thr-Ser)-Pro (Abcam, ab9344).

### In vitro kinase and phosphatase assays

His-tagged recombinant human PPARγ (Cayman) was attached to His-Tag dynabeads (Thermo Fisher) for 20 min. PPARγ-attached dynabeads were washed in kinase buffer (50 mM HEPES pH 7.5, 10 mM MgCl2) and resuspended in kinase buffer with 500 ng of ERK2 (ProQinase) or CDK5/p35 (Sigma-Aldrich) in the presence of 50 μM cold ATP and 0.15 μCi of γ(32P)ATP. Samples were incubated 5 min at 30°C. For in vitro de-phosphorylation, previously γ(32P)-ATP-phosphorylated PPARγ-attached beads were washed in phosphatase buffer (20 mM Tris, pH 7.5, 150 mM NaCl, 0.1% Triton X-100) and resuspended in phosphatase buffer with CDC14B or phosphatase-dead CDC14B. Samples were incubated 45 min at 30°C, and reactions were stopped by adding loading buffer and incubating 5 min at 95°C.

### Mass spectrometry

Proteins (∼5 μg) were reduced and alkylated by resuspending dried magnetic beads in 56 µL of 8M urea, 50 mM triethylammonium bicarbonate (TEAB, pH 8.5), 15 mM tris (2-carboxyethyl) phosphine (TCEP), and 50 mM chloroacetamide (CAA), followed by incubation for 1 hour at room temperature in the dark. Digestion was performed by adding 100 ng of Lys-C (Wako) and 100 ng of trypsin (Promega) per sample and incubating at 37°C for 16 hours. The resulting peptides were desalted using AttractSPE™ C18 disks (Affinisep), dried in a vacuum concentrator, and resuspended in 21 µL of 0.5% formic acid.

Liquid Peptide separation was performed using a Vanquish™ Neo HPLC system (Thermo Fisher Scientific) coupled to an Orbitrap Exploris 480 mass spectrometer (Thermo Fisher Scientific). Samples were first loaded onto a trap column (PepMap™ Neo, 5 µm C18, 300 µm × 5 mm) and subsequently separated on an Easy-Spray™ PepMap™ Neo analytical column (C18, 2 µm, 75 µm × 500 mm) maintained at 50°C. A 30-minute gradient was applied at a flow rate of 400 nL/min using buffer A (0.1% formic acid in water) and buffer B (100% acetonitrile with 0.1% formic acid). The gradient increased from 2% to 45% buffer B over 30 minutes, followed by a 4-minute wash at 98% buffer B. Peptides were ionized at 1.5 kV using the EASY-Spray source, with a capillary temperature of 300°C.

Samples were analyzed using two separate injections into the mass spectrometer. In the first injection, the instrument was operated in data-dependent acquisition (DDA) mode, alternating between MS and MS/MS scans using a Top 20 method. MS spectra were acquired over a mass range of 350–1400 m/z at a resolution of 60,000 FWHM (200 m/z), with an intensity threshold of 3.6 × 10⁵ and a dynamic exclusion of 15 seconds, excluding charge states of +1 and ≥ +6. Precursor ions were isolated using a 1.0 Th window and fragmented using higher-energy collisional dissociation (HCD) with a normalized collision energy of 29. MS/MS spectra were acquired at a resolution of 15,000 (200 m/z). The AGC target was set to 300% for MS scans with a maximum injection time (IT) of 25 ms, while for MS/MS scans, the AGC target was set to 100% with a maximum IT set to auto.

For the second injection, samples were analyzed in parallel reaction monitoring (PRM) mode to specifically identify and quantify peptides containing phosphorylation at S273. MS spectra were acquired over a mass range of 400–1370 m/z at a resolution of 60,000 FWHM (200 m/z). Ion peptides were isolated using a 1.0 Th window and fragmented using HCD with a normalized collision energy of 29. MS/MS spectra were acquired at a resolution of 60,000 (200 m/z). The AGC target values were set to 300% for MS scans and 250% for MS/MS scans, with a maximum IT of 25 ms for MS and 400 ms for MS/MS. The selected peptides for monitoring were TTDKS(ph)PFVIYDM(ox)NSLM(ox)M(ox)GEDK (m/z 850.6877, charge 3) and TTDKS(ph)PFVIYDMNSLM(ox)M(ox)GEDK (m/z 845.3561, charge 3).

Raw files acquired in DDA mode were processed with MaxQuant (v 1.6.0.16) using the standard settings against a Escherichia coli protein database (UniProtKB/TrEMBL, 4,305 sequences) containing the PPAR sequence and supplemented with contaminants. Carbamidomethylation of cysteines (+57.021 Da) was set as a fixed modification, whereas oxidation of methionines (+15.995 Da), phosphorylation of serine, threonine, and tyrosine (+79.966 Da), and protein N-terminal acetylation (+42.011 Da) were set as variable modifications. Minimal peptide length was set to 7 amino acids and a maximum of two tryptic missed-cleavages were allowed. Results were filtered at 1% FDR (peptide and protein level). Raw data were imported into Skyline (https://skyline.ms/). Label free quantification of identified phosphor-peptides was performed using the extracted ion chromatogram of the isotopic distribution. Only peaks without interference were used for quantification. Normalization was performed using the intensity of identified PPAR non modified peptides.

Raw files acquired in PRM mode were imported into Skyline and analyzed using a predicted spectral library from DeepPhospho^45^. Peptide identification and quantification required a dot product score of at least 0.84 for at least one sample^46^. The retention time at the apex of the quantified fragment ion peak group had to be within two minutes of the predicted retention time from DeepPhospho. S273-specific discriminant fragments were detected for all monitored peptide forms. Normalization was performed using the quantification of non-modified PPAR peptides from the DDA injections, along with the summed MS1 total ion current (TIC) from both DDA and PRM runs, to account for potential injection variability.

## Data availability

RNAseq data are available at the Gene Expression Ommibus with accession number GSE292631 (reviewer token: ozcfccaelfmfhoz). Proteomics data are available at ProteomeXchange Consortium via the PRIDE partner repository with the dataset identifier PXD062111.

## Acknowledgements

We thank Genomics, Bioinformatics and Protein Production Units for technical support. DVC was supported by a contract from Amigos CNIO. MS-R was supported by AECC (INVES18005SALA) and a Ramón y Cajal contract from the Ministry of Science, Innovation and Universities (RYC2020-028929-I). CVB was supported by contracts from Amigos CNIO, and the programs Juan de la Cierva and Ramón y Cajal (RYC2022-035259-I) from Agencia Estatal de Investigación, Ministry of Science, Innovation and Universities, MICIU. M.M. lab was supported by research grants from MICINN/AEI/FEDER (PID2021-128726, PDC2022-133408-I00 and PID2024-161681OB-I00), Comunidad de Madrid (Y2020/BIO-6519 and S2022/BMD-7437), Centro de Investigación Biomédica en Red de Cáncer (CIBERONC), and the iDIFFER network (RED2024-153635-T). VHIO would like to acknowledge the Cellex Foundation for providing research facilities and equipment and the CERCA Program from the Generalitat de Catalunya for their support on this research. CNIO (CEX2019-000891-S) and VHIO (CEX2020-001024-S/AEI/10.13039/501100011033) are Centers of Excellence Severo Ochoa (Agencia Estatal de Investigación, MICINN).

## Author Contributions

DCV, experimental work, data analysis and revision of the manuscript. AFM, experimental work and data analysis. AEB, experimental work in mice. OG and ASB, computational analysis. EZ and MI-proteomic analysis and revised the manuscript. MSR, experimental work and data analysis. CVB and MM supervised the project, analyzed data and wrote the manuscript.

