## Supplementary Figures and Tables for "CDC14 phosphatases control adipogenesis via PPARγ de-phosphorylation"

By Diana Vara-Ciruelos et al.

#### Supplementary Figures

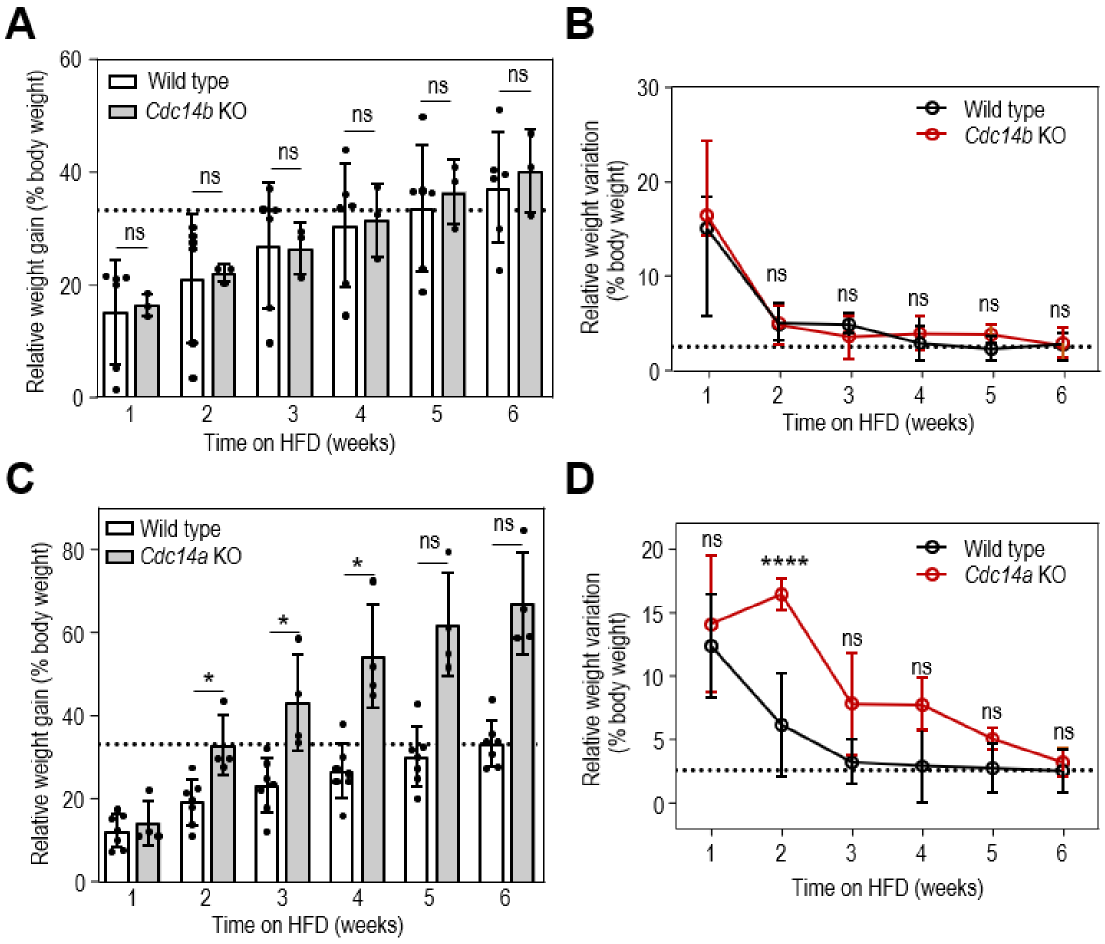


**Supplementary Figure 1. Different effect of *Cdc14a* and *Cdc14b* single depletion on weight gain under HFD. A, B.** Weight gain and weight variation respectively normalized to mouse individual body weight of wild type and *Cdc14b* KO mice throughout the 6 weeks of HFD treatment. Plots represent the represent the quantification of the weekly measurements during the experimental period. **C, D**. Weight gain and weight variation normalized to mouse individual body weight of wild type and Cdc14a KO mice throughout the 6 weeks of HFD treatment. Plots represent the represent the quantification of the weekly measurements during the experimental period.

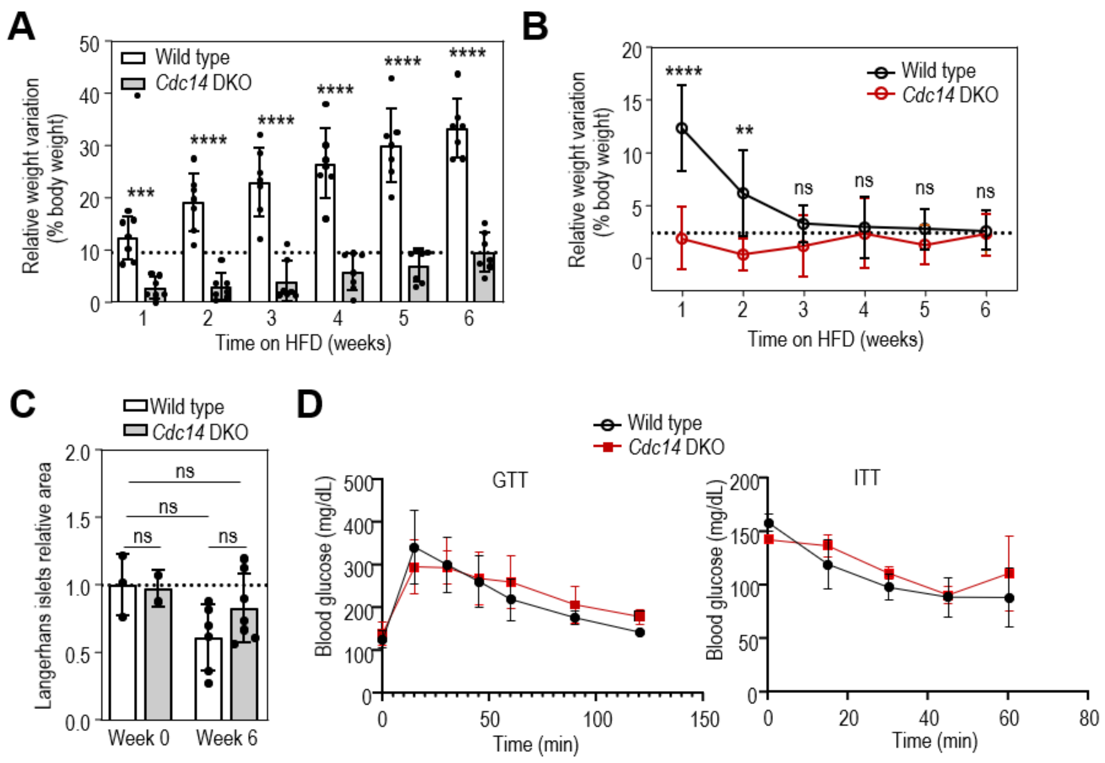


**Supplementary Figure 2. Defects in adipose tissue and glucose metabolism in *Cdc14a; Cdc14b*-mutant mice. A, B**. Weight gain and weight variation respectively normalized to mouse individual body weight of wild type and *Cdc14* DKO mice throughout the 6 weeks of HFD treatment. Plots represent the represent the quantification of the weekly measurements during the experimental period. **C**. Quantification of average area of pancreatic Langerhans islets in wild type and *Cdc14* DKO mice after 6 weeks of HFD or in control animals never subjected to HFD. Each dot represents the average relative area of at least 7 Langerhans islands per mice. **D**. Glucose tolerance test (GTT) and insulin tolerance test (ITT) of wild type and *Cdc14* DKO mice before HFD treatment. Data are shown as mean ± SEM. Statistical significance was assessed using the two-tailed Student's t-test with Welch correction. ns, not significant; *, p<0.05; **, p<0.01; ***, p<0.001; ****, p<0.0001.

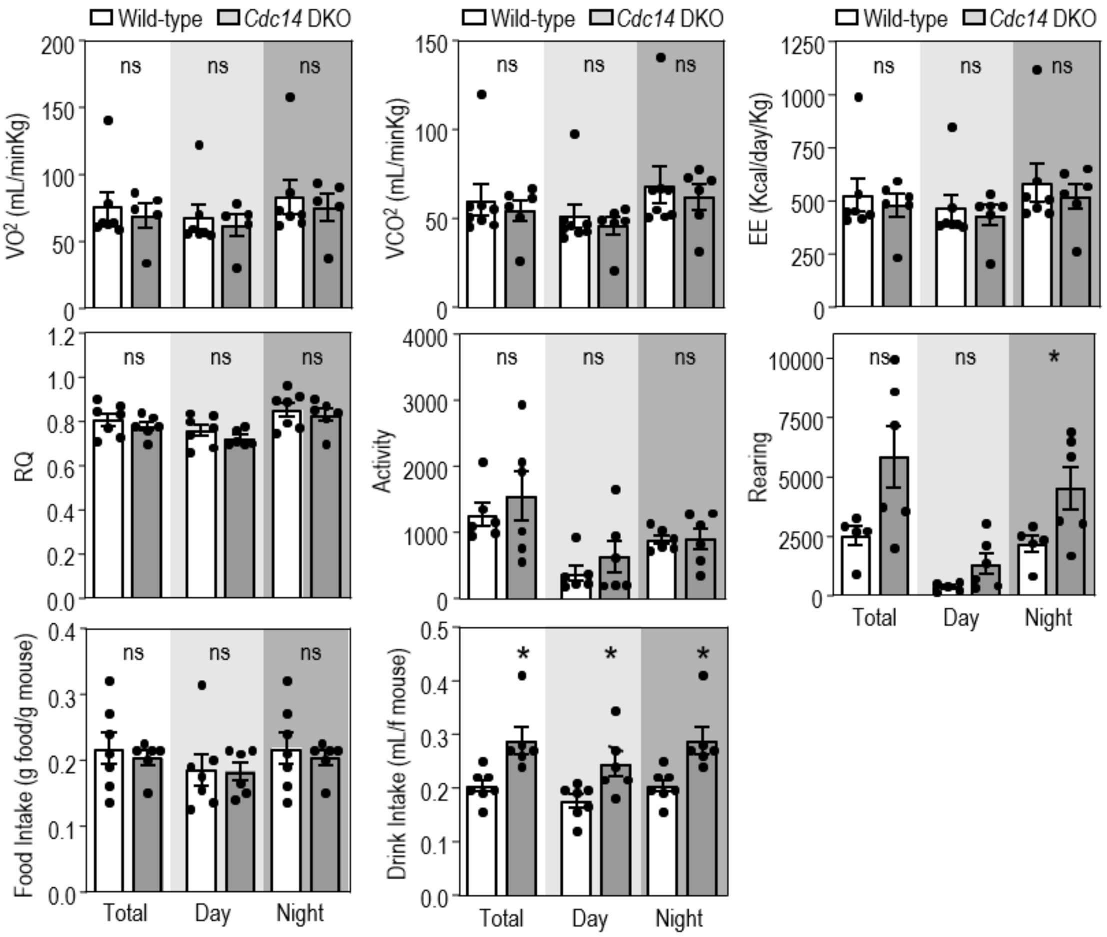


**Supplementary Figure 3. Cdc14 DKO mice exhibit normal energy expenditure.** No significant differences were found in oxygen consumption (VO2), carbon dioxide production (VCO2), energy expenditure (EE), respiratory quotient (RQ), global locomotive activity, rearing activity, food intake or drink intake between *Cdc14* DKO and control mice. Food intake and drink intake were measured throughout 72 hours during the dark (dark grey box) and light (light grey box) periods. Each point in all graphs represent the average measure of the corresponding parameter during the indicated period. N=7 wild type, N=6 *Cdc14* DKO. All mice used were males with an age comprised between 17 and 20 weeks. All data are normalized to the total body weight of each corresponding mouse. RQ = CO2 production/ Volume =2 consumption. EE is calculated using the semi-empirical Weir Equation. Data are shown as mean ± SEM. Statistical significance was assessed using the two-tailed Student's t-test with Welch correction. ns, not significant; *, $p<0.05$.

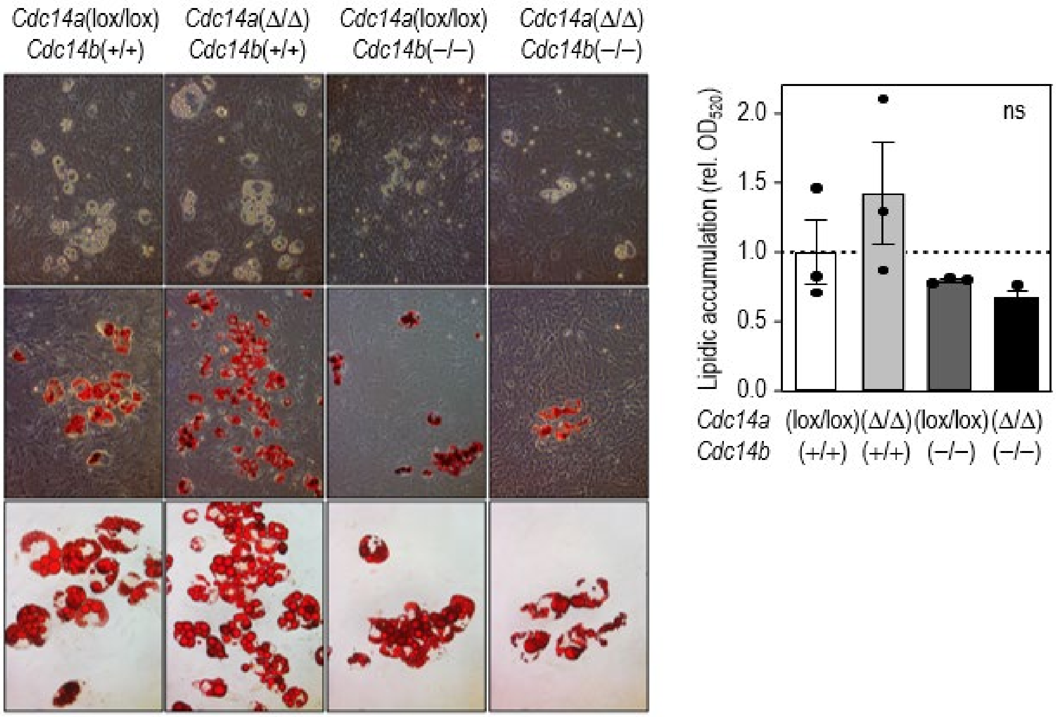


**Supplementary Figure 4. Title.** Representative images and quantification of Oil Red O staining in wild type, *Cdc14a* KO , *Cdc14b* KO and *Cdc14* DKO MEFs after 10 days of adipogenic differentiation. N=3 independent experiments. Data are means ± SD. Statistical significance was assessed using the two-tailed unpaired Student’s t-test with Welch correction. ns, not significant ($p>0.05$).

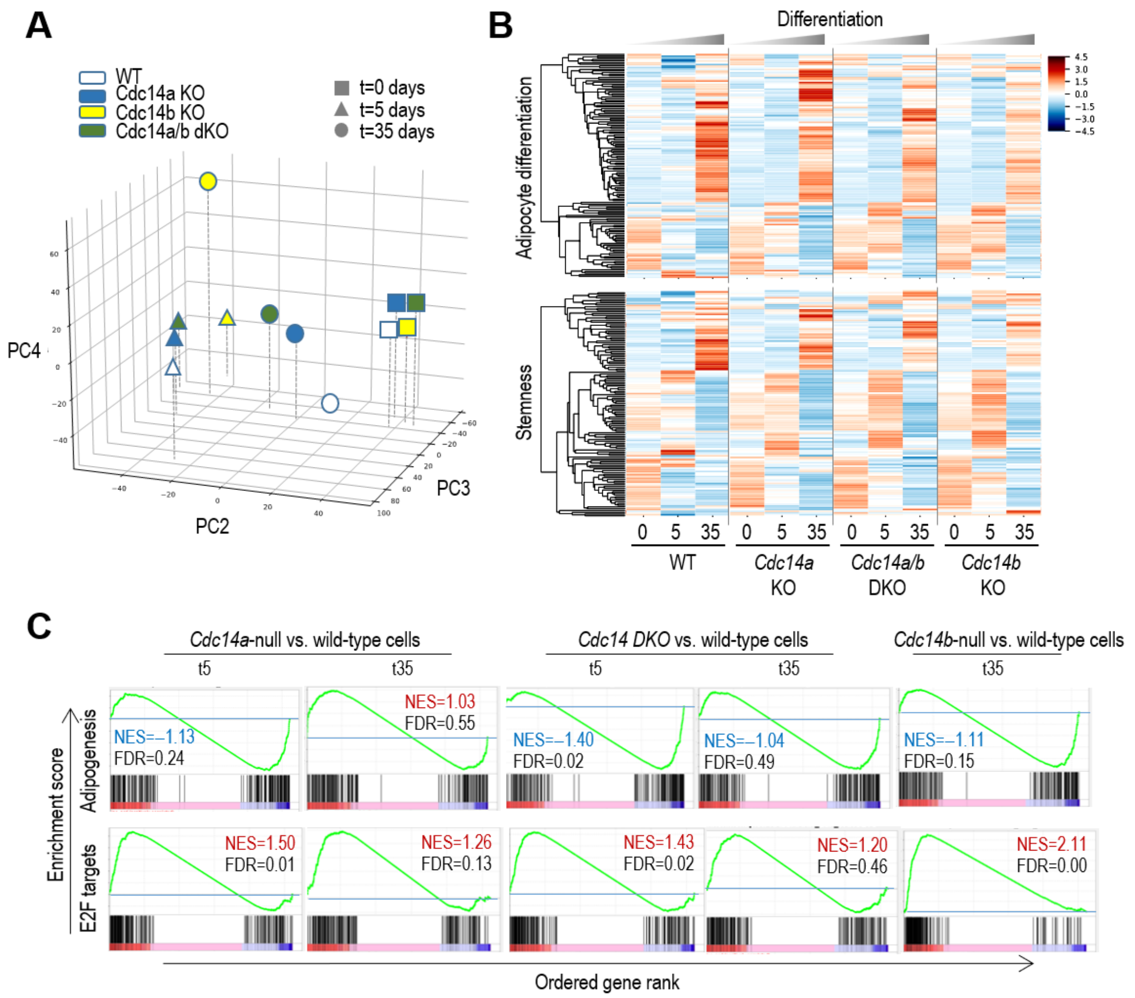


**Supplementary Figure 5. Transcriptional analysis of adipocyte differentiation from ESCs in the absence of CDC14. A**. Principal component analysis (PCA) of RNAseq data of mESCs wild type or depleted of CDC14A, CDC14B or CDC14A+B by CRISPR-Cas9 technology at days 0, 5 and 35 of the adipogenic differentiation protocol. **B**. Heat-map analysis of adipogenesis and stemness genes obtained by RNA-seq. **C**. Enrichment scores for the Hallmark signatures of adipogenesis and cell cycle transcripts (E2F targets) in the indicated conditions. The plots for Cdc14b-null cells at day 5 are shown in the min text (**Figure 6**).

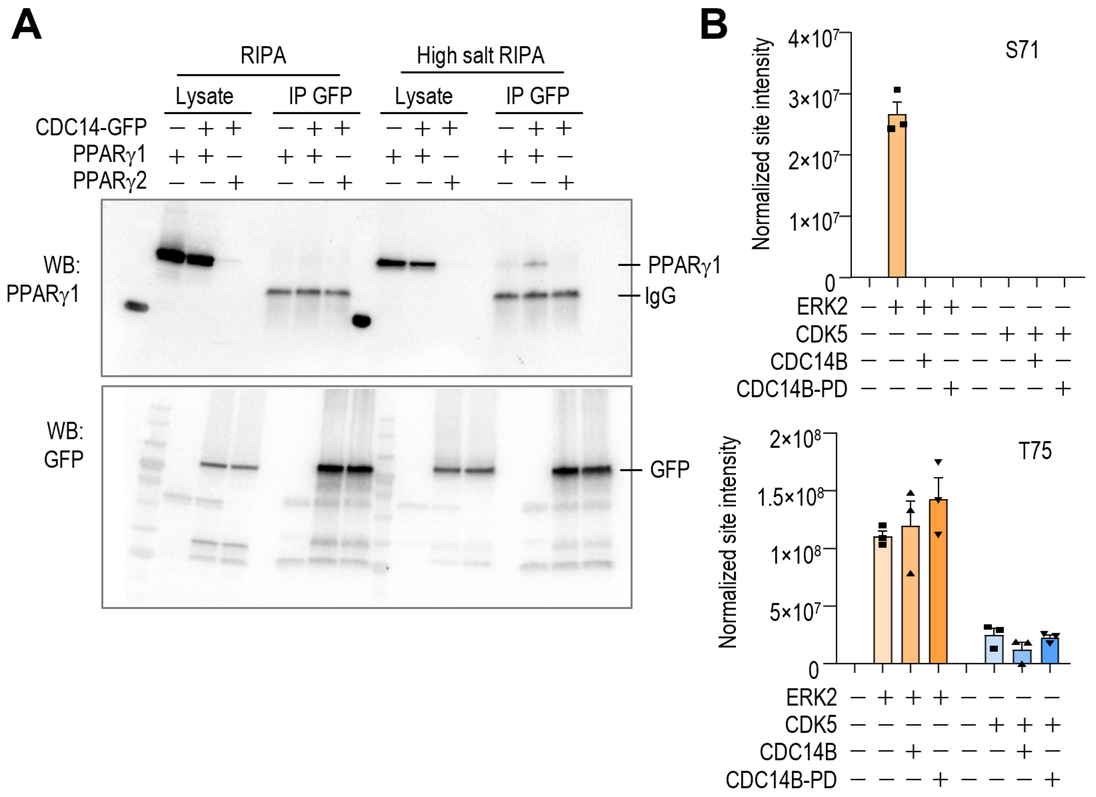


**Supplementary Figure 6. Interactions between CDC14B and PPARg. A**. Immunoblot analysis of CDC14B-GFP co-immunoprecipitations. HeLa cells were transfected with PPARγ 1 or 2 and CDC14B-GFP. Cytoplasmic and nuclear soluble proteins were extracted with RIPA, and nucleolar and chromatin-bound proteins were extracted with high salt RIPA. Immunoprecipitations with anti-GFP antibody were performed. Total protein extracts and immunoprecipitates were loaded and PPARγ (upper panel) and GFP (lower panel) were blotted. **B**. Phospho-proteomic analysis of ERK2 or CDK5-induced phospho-PPARγ2 after incubation with CDC14B or CDC14B-PD in T75 and S71 residues.

### Supplemental Tables

**Supplementary Table 1. Oligonucleotides used in this work.**

| Oligonucleotides | Forward | Reverse |
|---|---|---|
| **qPCR** | | |
| Cdc14a | GATCAGAATTTGCCGACCAG | ACCCACAATGATGCTTGTTTT |
| Cdc14b | GAAGCCTTCTCCAAACACCTT | CCACTGACTTCATCATCATCACT |
| Actin | GACGGCCAGGTCATCACTATTG | AGGAAGGCTGGAAAAGAGCC |
| **CRISPR sgRNA** | | |
| Cdc14a_sgRNA1 | CACCGACTACACCTCTTTCGACCAG | AAACCTGGTCGAAAGAGGTGTAGTC |
| Cdc14a_sgRNA2 | CACCGTCTTTTCCGCTGGTCGAAAG | AAACCTTTCGACCAGCGGAAAAGAC |
| Cdc14b_sgRNA1 | CACCGGTGGGCCCCATTCAAGATCC | AAACGGATCTTGAATGGGGCCCACC |
| Cdc14b_sgRNA2 | CACCGTTCCAGCCTGGATCTTGAAT | AAACATTCAAGATCCAGGCTGGAAC |
| **CRISPR t7** | | |
| Cdc14a | CCCTCCCCAACCCAAGAAAG | GACCAGCCTTGGTACTCAG |
| Cdc14b | CCAGGCTAATGACCCAAGCA | GAGCACACTTCTGCCTGGAT |
